# Proteome birthdating reveals age-selectivity of protein ubiquitination

**DOI:** 10.1101/2023.10.08.561433

**Authors:** Michael E. Meadow, Sarah Broas, Margaret Hoare, Fatemeh Alimohammadi, Kevin A. Welle, Kyle Swovick, Jennifer R. Hryhorenko, John C. Martinez, Seyed Ali Biashad, Andrei Seluanov, Vera Gorbunova, Abigail Buchwalter, Sina Ghaemmaghami

**Author notes:** Correspondence (SG).

## Abstract

Within a cell, proteins have distinct and highly variable half-lives. As a result, the molecular ages of proteins can range from seconds to years. How the age of a protein influences its environmental interactions is a largely unexplored area of biology. To investigate the age-selectivity of cellular pathways, we developed a methodology termed “proteome birthdating” that barcodes proteins based on their time of synthesis. We demonstrate that this approach provides accurate measurements of protein turnover kinetics without the requirement for multiple kinetic time points. As a first use case of the birthdated proteome, we investigated the age distribution of the human ubiquitinome. Our results indicate that the vast majority of ubiquitinated proteins in a cell consist of newly synthesized proteins and that these young proteins constitute the bulk of the degradative flux through the proteasome. Rapidly ubiquitinated nascent proteins are enriched in cytosolic subunits of large protein complexes. Conversely, proteins destined for the secretory pathway and vesicular transport have older ubiquitinated populations. Our data also identified a smaller subset of very old ubiquitinated cellular proteins that do not appear to be targeted to the proteasome for rapid degradation. Together, our data provide an age census of the human ubiquitinome and establish proteome birthdating as a robust methodology for investigating the protein age-selectivity of diverse cellular pathways.

**Significance Statement:** Cellular proteins have widely different ages - whereas some have been recently synthesized, others have existed in the cell for days or even years. How a protein’s age influences its functions and interactions is largely unknown because it is difficult to globally differentiate proteins based on their time of synthesis. To address this challenge, we developed an analytical method named “proteome birthdating” that can partition cellular proteins into multiple discernible age groups. As an example application, we used proteome birthdating to examine the protein age-selectivity of the ubiquitin proteasome system, a major protein degradation pathway in eukaryotes. Our results show that proteins destined for degradation by this pathway consist of either particularly young or particularly old proteins, with the former being the predominant population. Together, our results establish proteome birthdating as a useful approach for analyzing the turnover of proteins and investigating the functional consequences of their age.

## Introduction

Cellular proteins exist in a state of dynamic flux and are continuously synthesized and degraded (1). Within the proteome, turnover kinetics are highly variable and protein half-lives can therefore range from minutes to years (2–7). Turnover rates of proteins are often intimately linked to their function. For example, regulatory proteins typically have short half-lives in order for their steady-state levels to be rapidly responsive to changes in their synthesis rates. Conversely, abundant house-keeping proteins tend to be longer lived to minimize the energy expenditure associated with their continual synthesis and degradation (8). Because proteins are continuously turned over, the population of protein molecules within a cell varies widely in age. Whereas, some proteins have been very recently synthesized, others have persisted in the cell for days or even years (9). Investigating how a protein molecule’s age influences its modifications, interactions, and other properties within a cell is challenging as most biochemical experiments cannot differentiate proteins based on their time of synthesis.

An example of a protein modification that is likely to be age-selective is ubiquitination. It is known that a significant fraction of nascent proteins is ubiquitinated and targeted for proteasomal degradation during or shortly after translation (10–12). This process, carried out in part by the ribosome-associated quality control (RQC) pathway, is an important mechanism for clearance of translationally stalled or misfolded nascent proteins (13). Ubiquitin-mediated degradation of newly synthesized proteins also plays an important role in antigen presentation and the immune system by functioning as a source for MHC class I peptides (14). It is also known that when successfully synthesized proteins become damaged and aggregate over time, they are selectively targeted for ubiquitination and directed for degradation by the proteasome or selective autophagy (15–17). This process provides a mechanism for clearance of old, nonfunctional, or otherwise unneeded proteins (18). Given that ubiquitination appears to preferentially mark particularly young or particularly old proteins for degradation, the cellular ubiquitinome may be expected to have an age profile that is distinct from the proteome at large. However, to date, it has not been possible to analyze the age-selectivity of ubiquitination on proteome-wide scales.

A number of recently developed methods have combined time-resolved metabolic labeling with bottom-up proteomics to investigate protein dynamics on global scales (3, 19–22). These approaches, often referred to as “pulsed” or “dynamic” stable isotope labeling by amino acids in cell culture (dSILAC), typically employ a continuous labeling approach wherein an isotopically labelled precursor is introduced to cultured cells or whole model organisms over time. Changes in relative levels of newly synthesized proteins harboring the labeled precursor are measured in different experimental timepoints to determine the kinetics of protein turnover. These approaches have successfully measured protein half-lives on proteome-wide scales in many *in vitro* and *in vivo* model systems, providing fundamental insights into the post-transcriptional regulation of proteins.

In this study, we developed an alternative proteomic approach to dSILAC, named proteome birthdating, that utilizes a sequential rather than continuous labeling strategy. In proteome birthdating, a series of different isotopically labeled precursors are sequentially introduced to the same cell population over time and the relative level of each label is analyzed at a single experimental endpoint. By taking advantage of neutron encoded (NeuCode) amino acids (23, 24), we were able to barcode proteins with five labels, exceeding the multiplexing limits of typical SILAC experiments.

The application of proteome birthdating advances proteomic analyses in two distinct ways. First, it provides a methodology for analyzing protein half-lives that offers several advantages over dSILAC. Specifically, it allows proteome dynamics to be investigated by analyzing a single biological sample, rather than a series of experimental time points. Second, proteome birthdating provides an approach for partitioning proteins within a cell according to their molecular age. Thus, proteins can be distinguished in proteomic experiments based on their time of synthesis, and age-specific properties of proteins and cellular pathways can be investigated.

To demonstrate these two applications of proteome birthdating, we first used the approach to measure protein half-lives in primary human fibroblasts and compared the results to those obtained by traditional dSILAC. We then used the age-partitioned proteome to quantify the age distribution of the cellular ubiquitinome and investigate the age-selectivity of protein ubiquitination and proteasomal flux.

## Results

### Proteome birthdating by dynamic NeuCode labeling

The concept of proteome birthdating is illustrated in Figure 1A. In typical dSILAC experiments, isotopically labeled amino acids are introduced to a set of biological samples, each representing a distinct kinetic time point (3). Levels of fractional labeling are then measured within each sample at different time points to quantify protein turnover kinetics (Figure S1A). In contrast, in proteome birthdating, multiple labeled amino acids are sequentially added to, and then removed from, the same biological sample. Hence, each cell within the biological sample accumulates a mixture of differentially labeled proteins (Figure 1A, S1). In birthdated cells, relative levels of labels within proteins are determined by their turnover kinetics. For example, long-lived proteins will contain higher levels of labels introduced earlier in the time course, whereas short-lived proteins will contain higher levels of labels introduced later in the time course (Figure 1B, S1B). As such, end-point distributions of labels in a single sample can be used to determine proteome half-lives. Proteome birthdating not only enables analyses of proteome dynamics within a single biological sample, but it also provides a strategy for partitioning the proteome into experimentally discernible age groups. In birthdated cells, cellular proteins are demarcated based on their time of synthesis, allowing variations in age distributions of subsets of the proteome (e.g. differentially modified proteins) to be investigated by mass spectrometry (Figure 1C).

**Figure 1.**
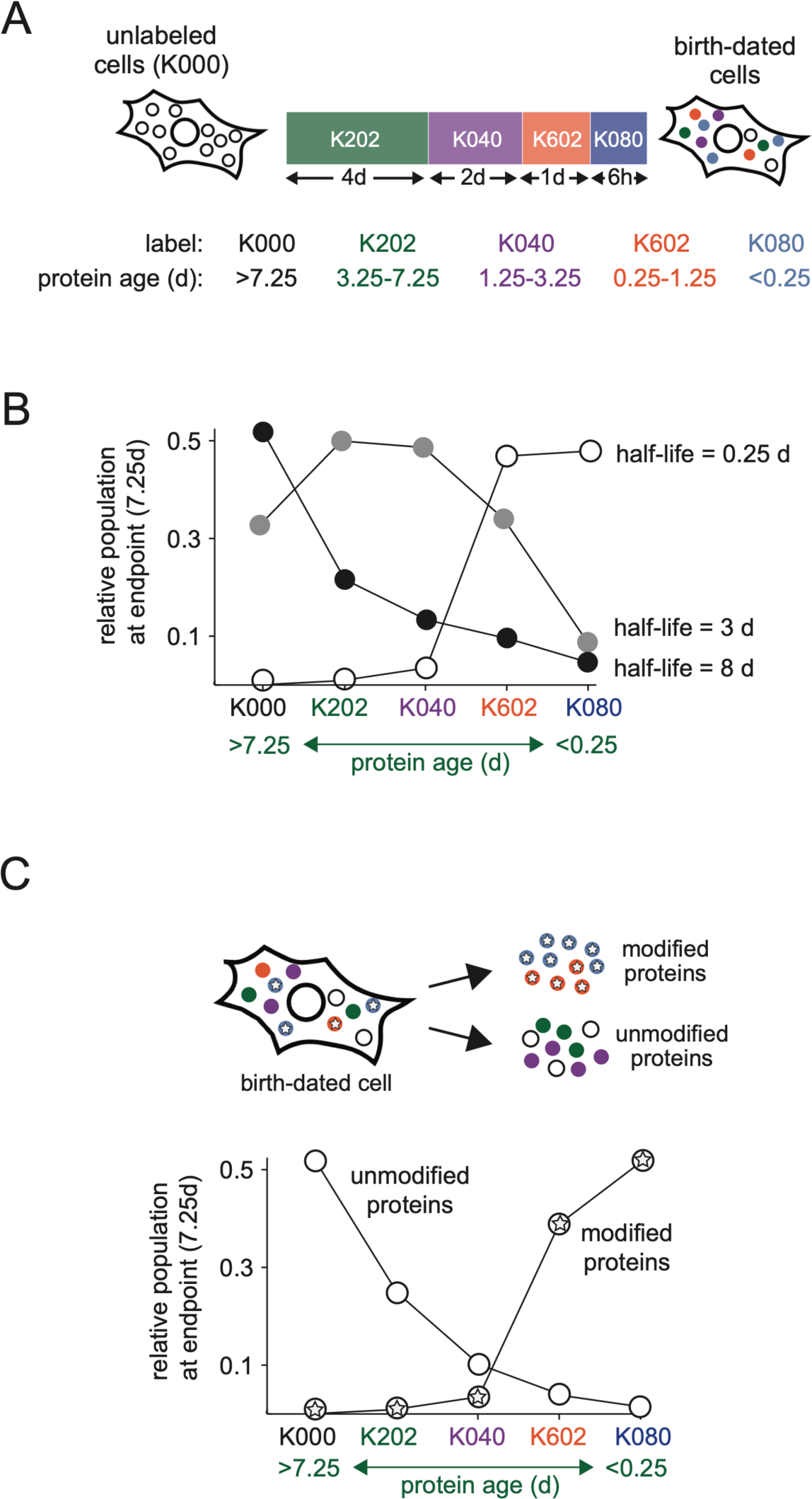
Proteome birthdating as a tool for measuring proteome turnover dynamics and age distributions. **A)** Experimental design of proteome birthdating in cell culture. Cells are sequentially labeled with isotopic variants of lysine (K_XYZ_) for the indicated lengths of time. The notation K_XYZ_, designates numbers of ^13^C (X), ^2^H (Y), and ^15^N (Z) isotopes. K_202_/ K_040_ and K_602_/ K_080_ are pairs of NeuCode amino acids. At the endpoint of the time course, protein molecules are differentially labeled with lysine variants in accordance with their age as listed in the table. **B)** The age distribution of a protein at the endpoint of the birthdating time course is dependent on its half-life. As examples, the plot shows K_XYZ_ distributions (i.e. age distributions) of three theoretical proteins with differing half-lives at the end of the labeling time course shown in (A). **C)** Proteome birthdating can be used as a tool to investigate the age selectivity of post-translational modifications (PTMs). The schematic shows a theoretical scenario where a PTM (star) preferentially modifies younger proteins. Also see Figure S1.

The proteome birthdating time course used in this study is illustrated in Figure 1A. Typically, the number of canonical SILAC metabolic labels that can be employed in an LC-MS/MS experiment is limited to three (commonly referred to as “light”, “medium” and “heavy”) (23, 25). This limitation is due to increased spectral complexities and reductions in proteome coverage that occurs upon incorporation of multiple SILAC labels. NeuCode amino acids circumvent this limitation by using isotopically labeled amino acids with mass differences in the order of milliDaltons (mDa) (23, 24). These slight differences in mass are generated using amino acid isotopologues that differ in distribution of neutrons within the molecule. Thus, NeuCode amino acids have the same nominal mass but vary slightly in exact mass because of mass defects caused by differences in nuclear binding energies. Differentially NeuCode labeled peptides behave as single spectral peaks in low resolution MS scans (and thus do not increase spectral complexities) yet can be resolved and quantified in high resolution scans. The use of NeuCode amino acids increases the number of labels that can be incorporated in birthdating experiments. In the experiments described in this study, five isotopologues of lysine, that include two pairs of NeuCode amino acids, are sequentially added to cells (Figure S1B). Lysine isotopologues are designated as K_XYZ_, where X, Y and Z respectively indicate numbers of ^13^C, ^2^H, or ^15^N isotopes in the molecule. The labeling time for each NeuCode label is sequentially decreased during the kinetic time course to capture the dynamics of proteins with a wide a range of half-lives.

### Measurement of protein half-lives in fibroblasts by proteome birthdating

We validated proteome birthdating by measuring protein half-lives in primary human dermal fibroblasts and comparing the results to those obtained by traditional dSILAC experiments. Turnover kinetics in human fibroblasts had also been analyzed by dSILAC in several previously published studies (8, 26–28). Here, fibroblasts were grown to quiescence and analyzed in parallel experiments by dSILAC over a labeling time course of one week, and by proteome birthdating using the labeling regime illustrated in Figure 1A. Experiments were conducted in non-dividing quiescent cells such that the labeling kinetics of proteins were solely dictated by their turnover and not cytosolic dilution resulting from cell division (28).

Spectra and labeling patterns for one example peptide ion are shown in Figure 2A. By proteome birthdating, we were able to analyze the labeling patterns of more than 50,000 peptide spectral matches (PSMs) mapped to more than 5,500 proteins in two biological replicate experiments (Figure S2, Supp. Data 1). Figure 2B illustrates age distributions of example proteins with either long (aldehyde dehydrogenase), intermediate (Chaperonin 60 - CPN60) or short (thrombospondin) half-lives. Protein half-lives were measured by fitting age distribution data to a first order kinetic model (see Materials and Methods). Half-life measurements for ∼3,750 proteins, determined in duplicate, passed designated quality control filters and were used in subsequent analyses. The range of half-lives for the covered human proteome spanned approximately two orders of magnitude (from ∼2 hours to ∼15 days), with a median measurement of 1.7 days (Figure 2C, Supp. Data 1). The overall age distribution of the proteome is shown in Figure 2D.

**Figure 2.**
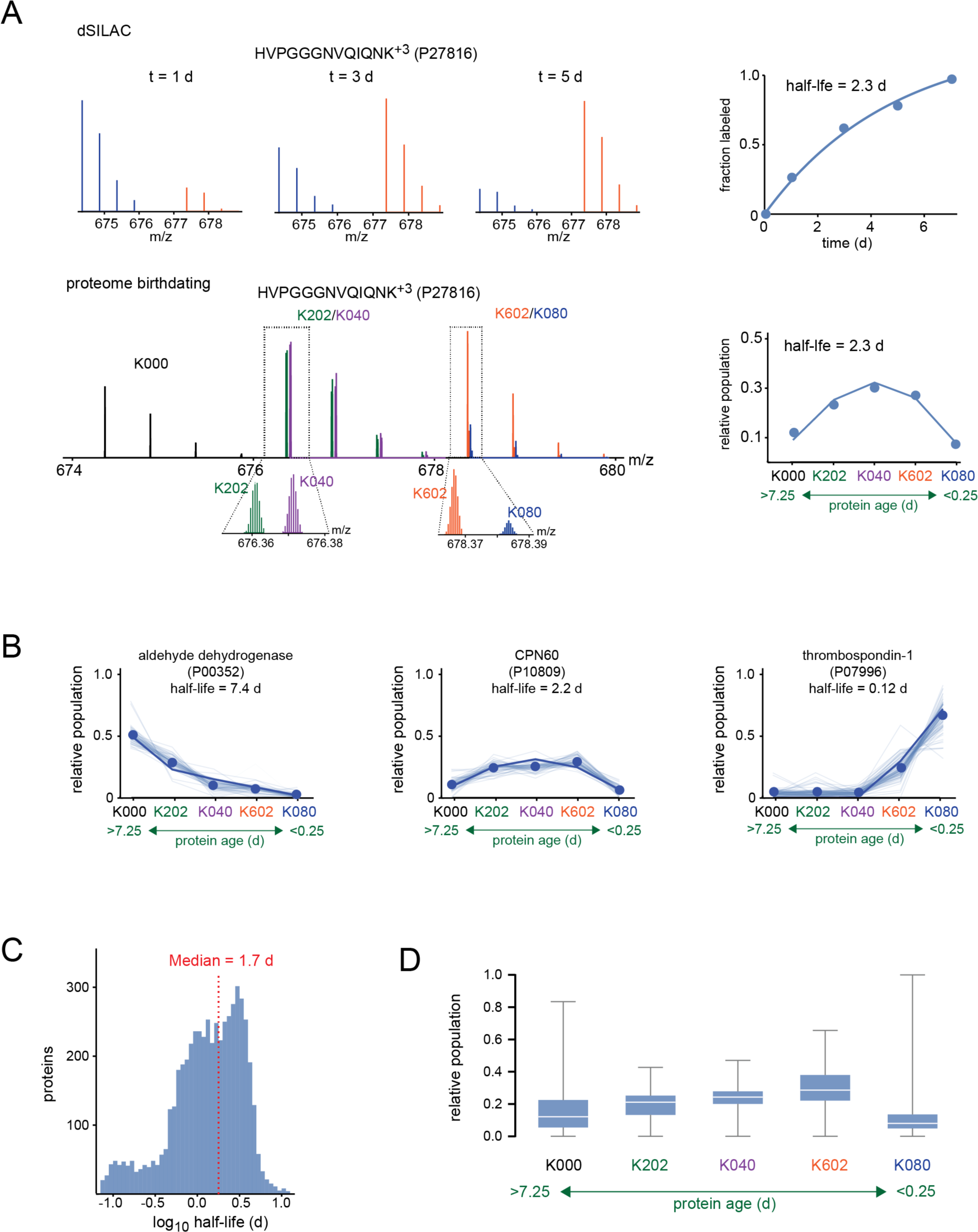
Global analysis of protein half-lives in human fibroblasts by proteome birthdating. **A)** Example spectral data and labeling kinetics of a peptide ion mapped to MAP4 protein (Uniprot: P27816, HVPGGGNVQIQNK^+3^) analyzed by dSILAC and proteome birthdating. In the dSILAC analysis, spectra were collected from distinct cultures exposed to a single isotopically labeled lysine (+6 Da) and arginine (+6 Da) for the indicated lengths of time. Unlabeled and labeled spectra are shown in blue and red colors, respectively. The half-life of the peptide was measured by fitting the time-resolved fractional labeling measurements to a first order kinetic model. In the proteome birthdating analysis, a single culture was sequentially labeled by multiple lysine variants as shown in Figure 1A. The spectrum and the plot show the relative intensities of each K_XYZ_-labeled form of the peptide at the end of the time course. The half-life of the peptide was measured by fitting the labeling pattern to a first order kinetic model (See Materials and methods) In the plots, dots are the measured data and blue lines indicate fit to the model. **B)** Example birthdating data for three proteins with either long (alcohol dehydrogenase, Uniprot: P00352), medium (CPN60, Uniprot: P10809), or short (thrombospondin 1, Uniprot: P07996) half-lives. Light blue lines indicate peptide-level data, dots indicate protein-level data (median of all measured peptides), and dark blue lines indicate fit to the model used to measure half-lives. The measured half-lives are shown above the plots. **C)** Distribution of protein half-lives in human fibroblasts measured by proteome birthdating. The dashed red line indicates the median half-life of the proteome (1.7 d). **D)** Box plots showing distributions of relative populations of each K_XYZ_ within the entire measured proteome. Boxes designate the interquartile range, white lines are the median, and whiskers indicate the entire distribution excluding far outliers (>2 SD from the mean). Also see Figure S2.

Several observations confirmed the precision and accuracy of half-life measurements determined by proteome birthdating. First, half-life measurements were reproducible in the two biologically replicated experiments (Figure S2B). Second, half-life measurements obtained by proteome birthdating correlated well to those obtained by canonical dSILAC in this study and previously published studies (26) (Figure S2C, D). Third, as would be expected, peptides mapped to the same proteins had similar measured half-lives (Figure S2E). Lastly, half-life measurements for subsets of the proteome expected to be composed of particularly short- or long-lived proteins largely had anticipated values. For example, signaling proteins and transcription factors had significantly shorter half-lives relative to housekeeping metabolic enzymes, components of the ribosome, histones, and nucleoporins that were previously determined to be very long-lived proteins (5, 9) (Figure S2F). Together, these data validated proteome birthdating as a robust methodology for measurement of protein half-lives that offers a number of practical and theoretical advantages over current dynamic proteomic approaches (see Discussion).

### Age distribution of human fibroblast ubiquitinome

We next sought to use proteome birthdating to investigate the age distribution of ubiquitinated proteins in human cells. Digestion of ubiquitinated proteins with trypsin generates peptides harboring characteristic di-glycine residues bound to lysine sidechains by isopeptide bonds (Kε-GG peptides). Kε-GG peptides can be readily enriched by selective antibodies and quantified by bottom-up proteomics (12, 29, 30). Typically, proteasome-bound polyubiquitinated proteins are transient and rapidly degraded after modification. To quantify the age distribution of a stabilized pool of ubiquitinated proteins, we first birthdated the fibroblast proteome and subsequently treated the cells with the proteasome inhibitor MG132 (Figure 3A). MG132-induced accumulation of ubiquitinated proteins was verified by Western blots (Figure S3A). In comparison to the experiments discussed in Figure 2, the birthdating time course was extended to 9.25 days to capture the dynamics of proteins with a wider range of half-lives. Following extraction and digestion, Kε-GG peptides were immunopurified and relative levels of NeuCode-labeled spectra were analyzed by LC-MS/MS as described above. In all, we were able to characterize the age distribution of 1,846 Kε-GG peptides mapped to 1,089 proteins (Figure S3B, Supp. Data 2). For comparison, unmodified peptides were similarly generated from both control cells and cells treated with MG132 and used to quantify the age distribution of unmodified proteins.

**Figure 3.**
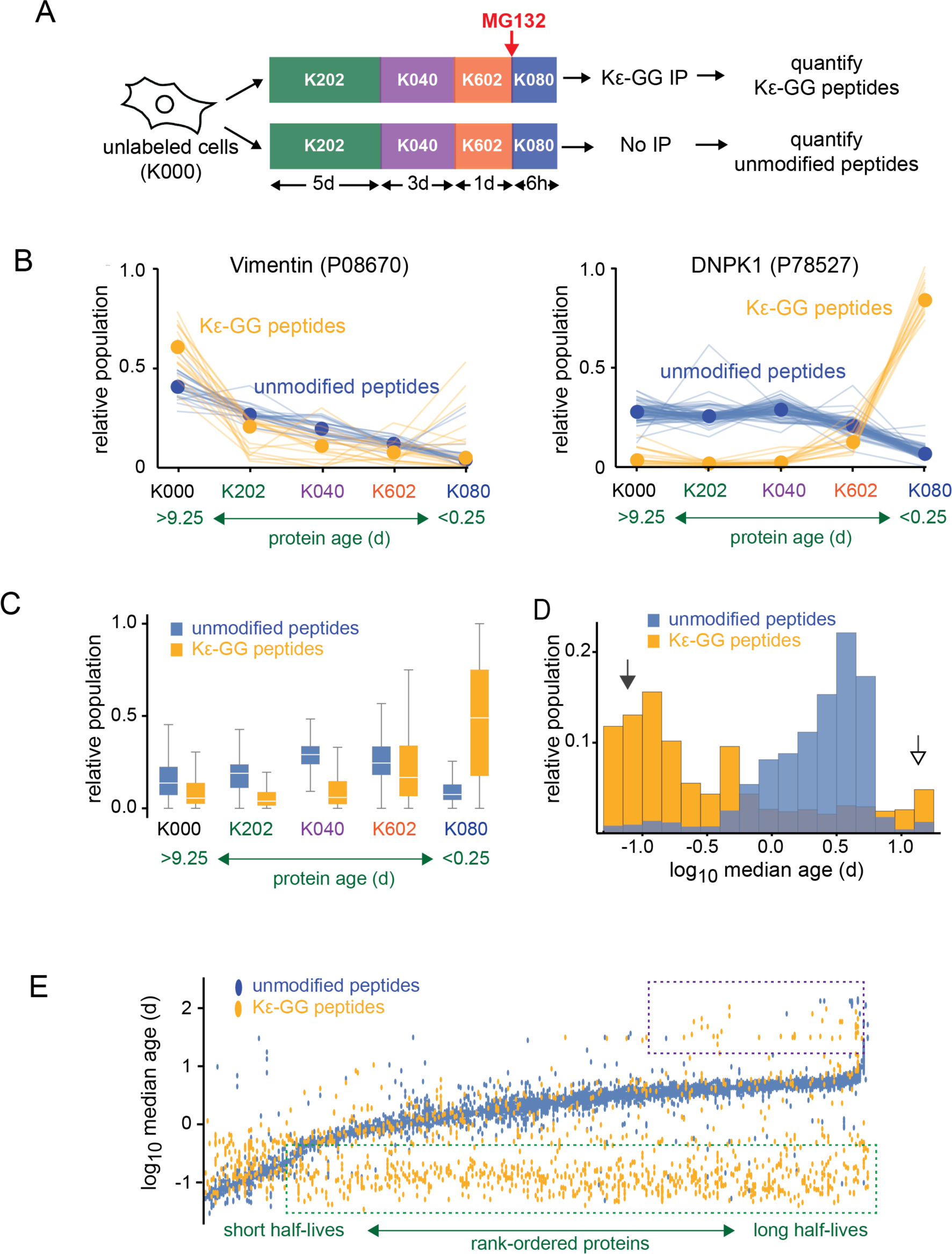
Age distribution of Kε-GG peptides in human fibroblasts. **A)** Experimental design for birthdating of Kε-GG peptides. The red arrow indicates the point of addition of the proteasome inhibitor MG132. In cells treated with MG132, Kε-GG peptides were enriched by immunopurification prior to MS analysis. **B)** Birthdating data for Kε-GG and unmodified peptides mapped to two example proteins: vimentin (Uniprot: P08670) and DNPK1 (Uniprot: P78527). Faint lines indicate peptide-level data and solid lines and dots indicate median of all measured peptides for the protein. Yellow and blue colors designate Kε-GG and unmodified peptides, respectively. **C)** Distributions of relative populations of K_XYZ_ labels within all measured Kε-GG and unmodified peptides. Box plots were constructed as described in Figure 2D. **D)** Distribution of median ages of Kε-GG and unmodified peptides. Close and open arrows respectively highlight the preponderance of Kε-GG peptides that are particularly young and old relative to the age distribution of unmodified peptides. **E)** Comparison of median ages of Kε-GG and unmodified peptides mapped to the same proteins. Vertical columns of points represent data for peptides matched to specific proteins that are rank ordered based on their (unmodified) half-lives. Green and purple dashed boxes indicate Kε-GG peptides that are, respectively younger and older than their unmodified counterparts mapped to the same proteins. Also see Figure S3.

In general, age distributions of Kε-GG and unmodified peptides mapped to the same protein differed markedly from each other (Figure 3B). For some proteins (e.g., vimentin shown in Figure 3B), Kε-GG peptides were significantly older than unmodified peptides. However, most Kε-GG peptides were significantly younger than their unmodified counterparts in the same protein (e.g., DNPK1 in Figure 3B.) The global age distributions of Kε-GG and unmodified peptides are shown in Figure 3C. Most Kε-GG peptides harbored the K080 label, indicating that they were synthesized in the cell within the last 6 hours of the experimental time course. The second most frequent label in Kε-GG peptides was K000, marker of the oldest group of proteins, which were synthesized more than 9 days prior to proteasomal inhibition.

Median ages of Kε-GG and unmodified peptides were measured based on the assumption that both are derived from proteins that display first order turnover kinetics (see Materials and Methods). Figure 3D compares the median age distributions of all Kε-GG and unmodified peptides. The data indicated that globally, Kε-GG peptides tended to be either much younger or much older than unmodified peptides, with the younger modified peptides being the overall more prevalent form (arrows in Figure 3D). Unlike unmodified peptides, Kε-GG peptides mapped to the same protein generally had a broad range of ages, suggesting that modifications of different sites within the same protein had distinct age-selectivity (Figure 3E). For some long-lived proteins, Kε-GG peptides appeared to be very old, with median ages exceeding 30 days (Figure 3E, S3C, dashed purple box). However, most Kε-GG peptides were significantly younger than unmodified peptides mapped to the same protein (Figure 3E, S3C, dashed green box). Young Kε-GG peptides were derived from both short-lived and long-lived proteins. In fact, many Kε-GG peptides that were only a few hours old mapped to very stable proteins whose steady-state half-lives were more than 10 days (Figure 3E, S3C, D).

### Correlation between protein age and proteasomal flux

Kε-GG peptides can be generated by polyubiquitinated proteins targeted to the proteasome, as well as proteins modified by ubiquitin or ubiquitin-like proteins (e.g. NEDD8 and ISG15) that are not necessarily marked for proteasomal degradation (31). To differentiate Kε-GG-generating proteins based on their contribution to proteasomal flux, we measured changes in steady-state levels of Kε-GG peptides after proteasomal inhibition. Fibroblasts were treated with MG132 for 6 hours and changes in steady state levels of Kε-GG and unmodified peptides were analyzed by LC-MS/MS (Figure 4A, S4A). As had been observed in previous studies (12, 29, 30), levels of some, but not all, Kε-GG peptides increased upon proteasomal inhibition (Figure 4B, Supp. Data 3). Conversely, levels of non-Kε-GG peptides were not impacted by the addition of MG132 (Figure S4B).

**Figure 4.**
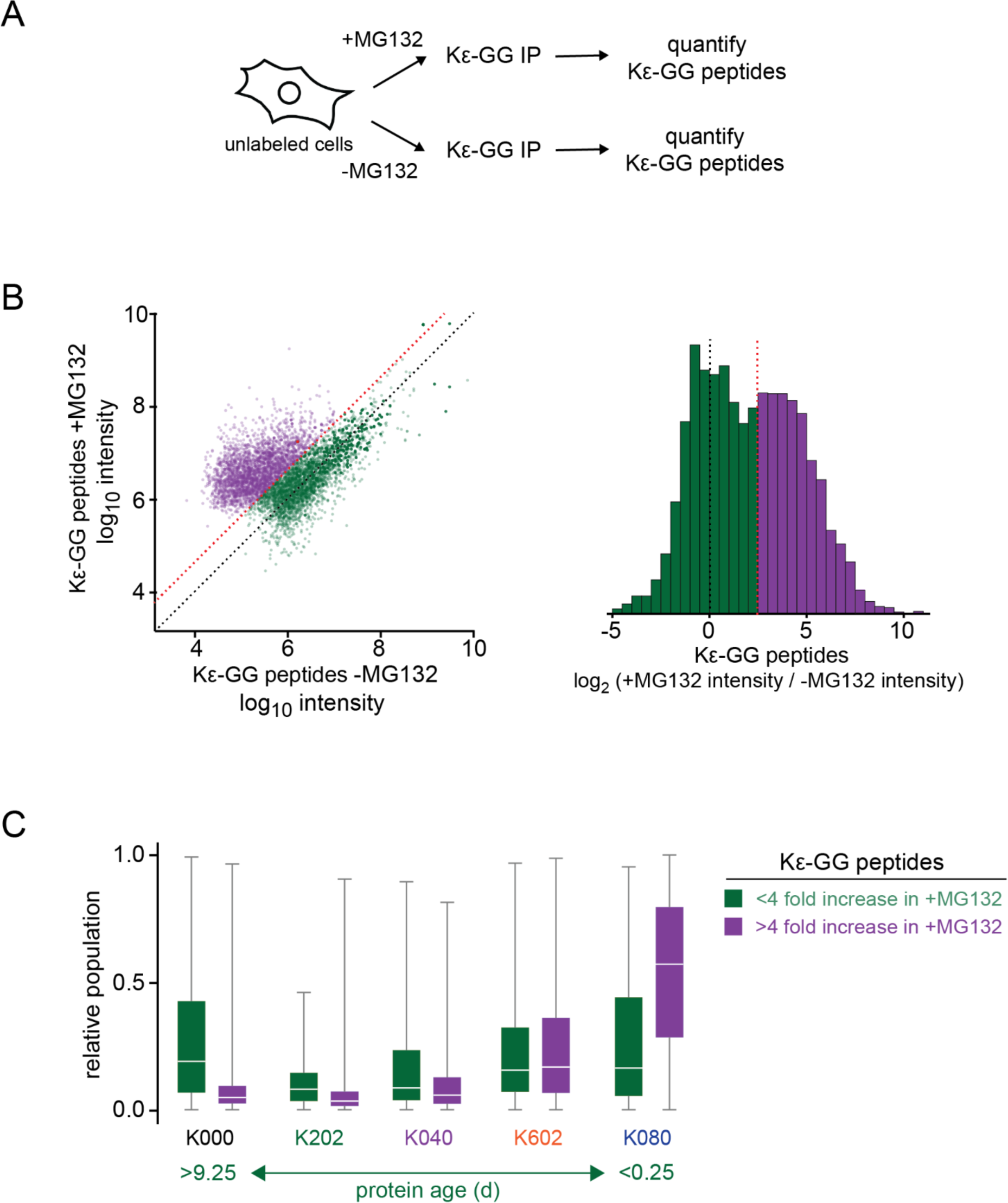
Relationship between age and proteasomal flux of Kε-GG peptides in human fibroblasts. **A)** Experimental design for measurement of changes in Kε-GG peptide levels upon proteasomal inhibition. **B)** Left plot shows the pairwise comparison of Kε-GG peptide intensities in the presence and absence of MG132. Peptides whose levels increased by greater or less than a factor of four are shown in purple and green colors, respectively, and are separated by the dashed red line. The dashed black line is the identity line. The histogram to the right represents the log_2_ intensity ratios of Kε-GG peptides in the presence and absence of MG132. **C)** Comparison of age distributions of Kε-GG peptides determined by proteome birthdating as measured in Figure 2. The data indicate that Kε-GG peptides that are rapidly targeted to the proteasome are significantly younger than those that are not rapidly targeted to the proteasome. The box plots were constructed as described in Figure 2D.

Kε-GG peptides mapped to the same protein generally accumulated at variable levels upon proteasomal inhibition (Figure S4C). Thus, Kε-GG-generating modifications at different sites within the same protein targeted proteins for proteasomal degradation at variable efficiencies. Analysis of Kε-GG peptides mapped to ubiquitin itself indicated increases in K6, K11, K27, K33, and K48 ubiquitin-ubiquitin linkages in MG132 treated cells, with the most dramatic increase observed in K48 linkages (Figure S4D). Conversely, levels of K63 linkages remained constant. These results are consistent with previous studies indicating that K63 linked polyubiquitin linkages do not target proteins for proteasomal degradation (12, 29).

We next compared age distributions of Kε-GG peptides to changes in their expression levels following proteasomal inhibition. Our analysis indicated that MG132-induced changes in levels of Kε-GG peptides were inversely correlated to their age (Figure S4E, F). To further analyze this trend, we compiled a list of ∼9,000 Kε-GG peptides mapped to ∼2,600 proteins that increase their levels by greater than 4-fold upon MG132 treatment (Figure 4B purple, Supp. Data 3). This subset of Kε-GG peptides can be inferred to represent a subset of the ubiquitinome that is rapidly targeted to the proteasome. Figure 4C compares the age distributions of the proteasome-targeted subset of Kε-GG peptides to those that did not significantly accumulate upon proteasomal inhibition. The data indicate that targets of the proteasome were enriched in the youngest subset of Kε-GG peptides, whereas older Kε-GG peptides were relatively minor contributors to overall proteasomal flux. Thus, most proteasome-bound proteins that accumulated after 6 hours of MG132 treatment were nascent polypeptides that were modified during or within a few hours after their synthesis. Conversely, proteins that were modified days after synthesis were lesser contributors to proteasomal flux.

### Gene ontology (GO) enrichment analysis of young and old proteasomal targets

We next searched for GO annotations that were statistically enriched in particularly young or particularly old proteasomal targets. For each identified proteasomal target, median ages of Kε-GG peptides were measured relative to unmodified peptides (Figure S3D, Supp Data 2.) GO annotations whose constituent proteins had median Kε-GG peptide ages differing significantly from the overall population median were identified (Supp Data 2). Among the ontologies with the youngest relative Kε-GG peptides were large stoichiometric complexes, including the 20S proteasome, Arp2/3 complex, COP9 signalosome, CTT chaperonin complex, the nucleopore complex and the WASH complex (Figure 5A). Relative to the rest of the proteome, proteins known to be subunits of protein complexes generated younger Kε-GG peptides targeted to the proteasome. However, complex subunits were not solely responsible for generating proteasome-bound young Kε-GG peptides. Even among proteasome targets that are known to be monomeric, Kε-GG peptides were generally younger than their unmodified counterparts mapped to the same proteins.

**Figure 5.**
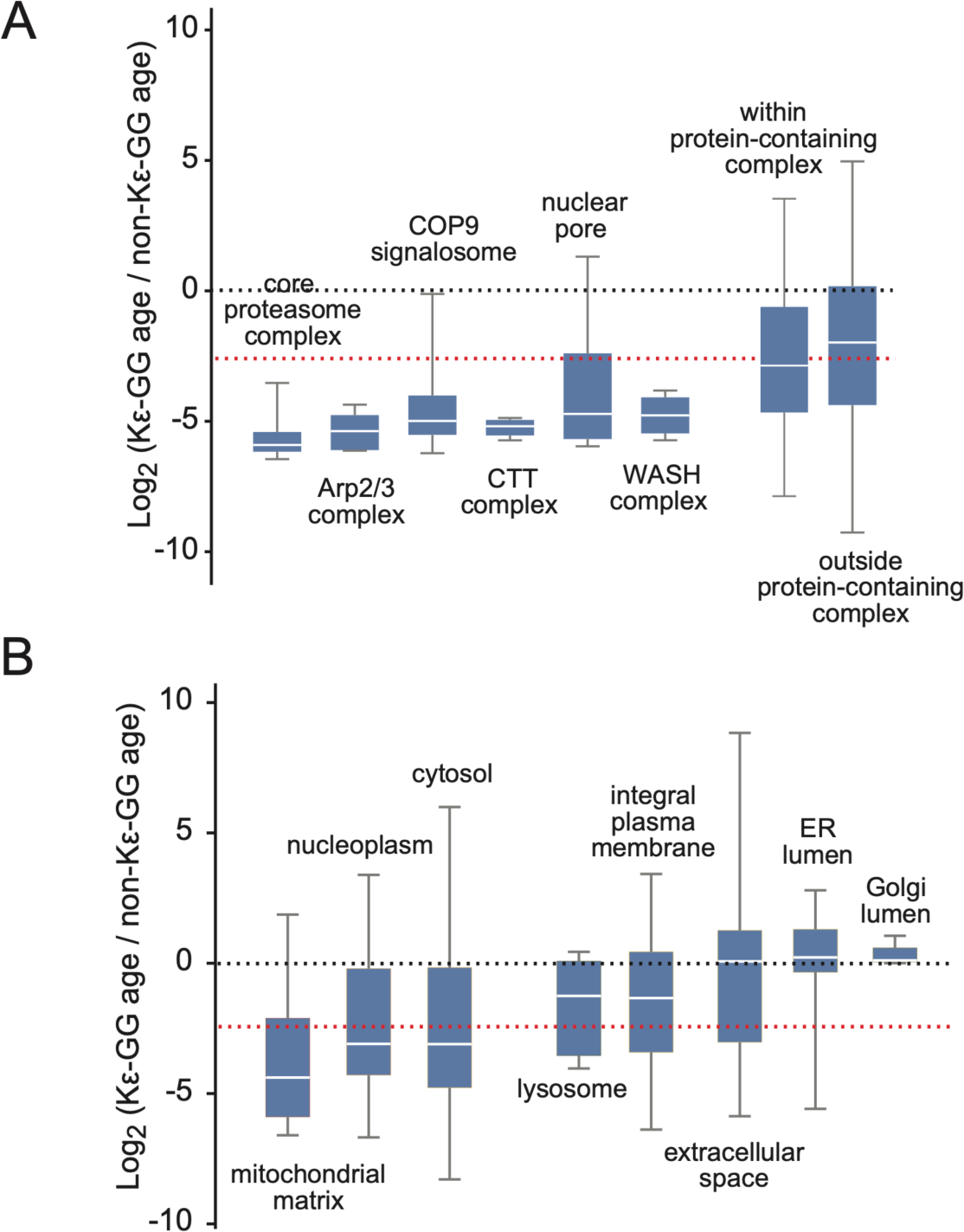
Age distributions of proteasomally targeted Kε-GG peptides mapped to gene ontologies (GO). **A)** Age distributions of Kε-GG peptides relative to unmodified peptides for proteins mapped to GO terms associated with specific protein complexes. “Within protein-containing complex” includes all proteins mapped to terms of GO:0032991 (“protein-containing complex”), and “outside protein-containing complex” includes all proteins excluded from this set. The dashed red line indicates the mean value for age distributions of all Kε-GG peptides relative to unmodified peptides for the entire proteome. **B)** Age distributions of Kε-GG peptides relative to unmodified peptides for proteins mapped to GO terms associated with major subcellular localizations. The complete list of GO terms with Kε-GG peptide age distributions that significantly deviate from the proteome mean is included in Supplementary Table S2.

GO enrichment analyses also identified significant variations in age distributions of proteasomal targets mapped to proteins localized to different subcellular compartments. Specifically, proteins localized to the cytosol, nucleus and mitochondria had younger proteasomal targets in comparison to those localized to plasma and organelle membranes, extracellular space, and lumens of the lysosome, ER, and Golgi apparatus (Figure 5B). The results indicate that rapid degradation following synthesis is less prevalent among nascent proteins that are co-translationally targeted for the secretory pathway or vesicular transport to organelles.

## Discussion

We conducted experiments in human fibroblasts to validate and demonstrate the utility of a dynamic proteomic approach termed proteome birthdating. In proteome birthdating, a single biological sample is sequentially labeled with multiple isotopically labeled amino acids for varying time intervals. Birthdated cells amass proteins that are differentially labeled in accordance with their times of synthesis (i.e. “age”). We show that proteome birthdating is useful in at least two types of analyses. First, it provides a methodology for measuring protein half-lives on a global scale. Second, it can determine the age profiles of specific subsets of the proteome.

In proof-of-concept studies, we took advantage of NeuCode amino acids that are commercially available to partition the proteome into five distinct age groups. This level of multiplexing surpasses the typical limit of three metabolic labels imposed by standard SILAC experiments. However, the number of NeuCode labels potentially resolvable by modern mass spectrometers expands well beyond five. For example, tryptic peptide ions harboring at least 8 different NeuCode variants of the +8 lysine isotopolgue are resolvable at resolutions exceeding 500,000 (a capability that is available in many current mass analyzers), and the level of multiplexing can be further expanded by inclusion of other lysine and arginine isotopologues (23). Thus, with the synthesis of additional NeuCode amino acids, the time resolution of proteome birthdating experiments may be greatly expanded, allowing proteomes to be partitioned into larger numbers of age ranges.

As a global method for measuring protein half-lives, proteome birthdating offers several advantages over canonical dSILAC continuous labeling strategies. In dSILAC, each kinetic timepoint is represented by a distinct biological sample. In contrast, in proteome birthdating, a single multi-labeled sample provides sufficient temporal information to accurately discern protein half-lives. This feature reduces the size and cost of dynamic proteomic projects in cell culture and complex organisms and may facilitate analyses of larger sample sizes. Additionally, as complete labeling time courses are encompassed within a single biological sample, signal intensities of all isotopic labels are internally normalized and experimental errors associated with rate constant measurements based on multiple time points are reduced. Importantly, distributions of multiple temporal labels within each birthdated cell allows for accurate determination of protein half-lives within that individual cell. With recent advances in mass spectrometry instrumentation making single-cell proteomics more commonplace (32–34), proteome birthdating may open the door to investigations of multi-timepoint proteome-wide protein turnover kinetics within individual cells.

Cellular proteomes undergo continuous cycles of synthesis and degradation at rates that vary by orders of magnitude. As a result, within cells, ages of individual protein molecules are highly variable. With a few exceptions (35, 36), the relationship between protein molecular age and function remains largely unexplored. Proteome birthdating provides a methodology for investigating age distributions of specific subsets of the proteome and enables analyses of protein age-selectivity of cellular pathways. As an example of this type of application, we investigated the age distribution of the human ubiquitinome.

It has been shown that the UPS is responsible for degradation of aberrant ribosome-bound and nascent proteins, removal of old and misfolded proteins, and basal turnover of specific short-lived proteins (10–18, 37). However, to the best of our knowledge, relative contributions of each of these target sets to total proteasomal flux had remained unquantified. Our data indicate that the overall age distribution of the proteasome-bound ubiquitinome is significantly younger than the steady state population of cellular proteins. Based on this observation, we conclude that newly synthesized proteins are the majority constituents of the population of ubiquitinated proteins destined for degradation by the proteasome.

Young proteins can be ubiquitinated and targeted to the proteasome even in cases where the steady-state population of that protein in the cell is relatively long-lived. The results support previous findings indicating that many proteins are rapidly culled during or soon after synthesis, and those that survive become part of a more stable steady-state proteome (35, 38). Functional enrichment analysis identified subunits of protein complexes as one common class of rapidly degraded proteins. This result suggests that newly synthesized subunits that fail to incorporate into their cognate complexes are targeted for rapid degradation by the UPS. However, our data indicate that degradation of nascent polypeptides is not a unique property of protein complex subunits and appears to be prevalent across much of the proteome. Interestingly, the ubiquitinome of proteins that are known to be initially released into the cytosol following translation (e.g. proteins ultimately localized to the cytosol, nucleus, or mitochondrion) were generally younger than those destined for the secretory or vesicular transport pathways. This result may indicate that co-translational interactions of nascent proteins with the signal recognition particle (SRP) and their transport to the ER (39) may act as a protective mechanism or delay the ubiquitination and eventual degradation of misfolded newly synthesized proteins. However, additional experiments are needed to uncover the mechanistic basis of this effect.

Our data identified a minor subset of the ubiquitinome that was composed of relatively old proteins. However, cellular levels of these modified proteoforms did not significantly increase upon proteasomal inhibition, indicating that they are relatively minor contributors to overall proteasomal flux. We propose three non-mutually exclusive interpretations of this observation. First, ubiquitin or ubiquitin-like adducts of some older proteins may represent non-degradative or stabilizing modifications (40). Second, although older ubiquitinated proteins may be targeted to the proteasome, their degradation may be occurring more slowly than younger ubiquitinated proteins. For example, older proteins may be enriched in misfolded conformers whose degradation is impaired due to aggregation (41). Third, older misfolded or aggregated proteins that are ubiquitinated may be targeted for degradation by non-proteasomal pathways. For example, ubiquitin-dependent autophagic pathways such as selective aggrephagy may play a predominant role in clearance of older misfolded proteins (42).

A model consistent with our observations is presented in Figure 6. In this model, a fraction of newly synthesized proteins is ubiquitinated and rapidly targeted for proteasomal clearance. This population of young ubiquitinated proteins is the dominant contributor to overall proteasomal flux in the cell. As has been previously shown, UPS-mediated degradation of newly synthesized proteins represents an important quality control mechanism to clear proteins that fail to successfully undergo translation, fold, or incorporate into their cognate protein complexes, and may play an important role in MHC 1 antigen presentation (10–15). Nascent proteins that survive immediate proteolysis become part of the cell’s accumulated protein population. The clearance of this steady-state population occurs largely according to non-selective (random) first order kinetics. As proteins get older and accumulate structural damage, they again can become targets of age-selective ubiquitination and targeted for degradation by the ubiquitin-dependent pathways such as the UPS or selective autophagy. However, selectively targeted younger and older populations of proteins are generally not present at high levels relative to their accumulated steady-state cellular populations. This is because targeted younger proteins are rapidly cleared before contributing to the steady state proteome, and older proteins generally do not become targets of selective degradation until their age significantly surpasses their basal half-life, a feat accomplished by a small fraction of their population. Thus, at steady state, the majority of a protein’s accumulated population appears to be degraded randomly with a uniform first order rate constant.

**Figure 6.**
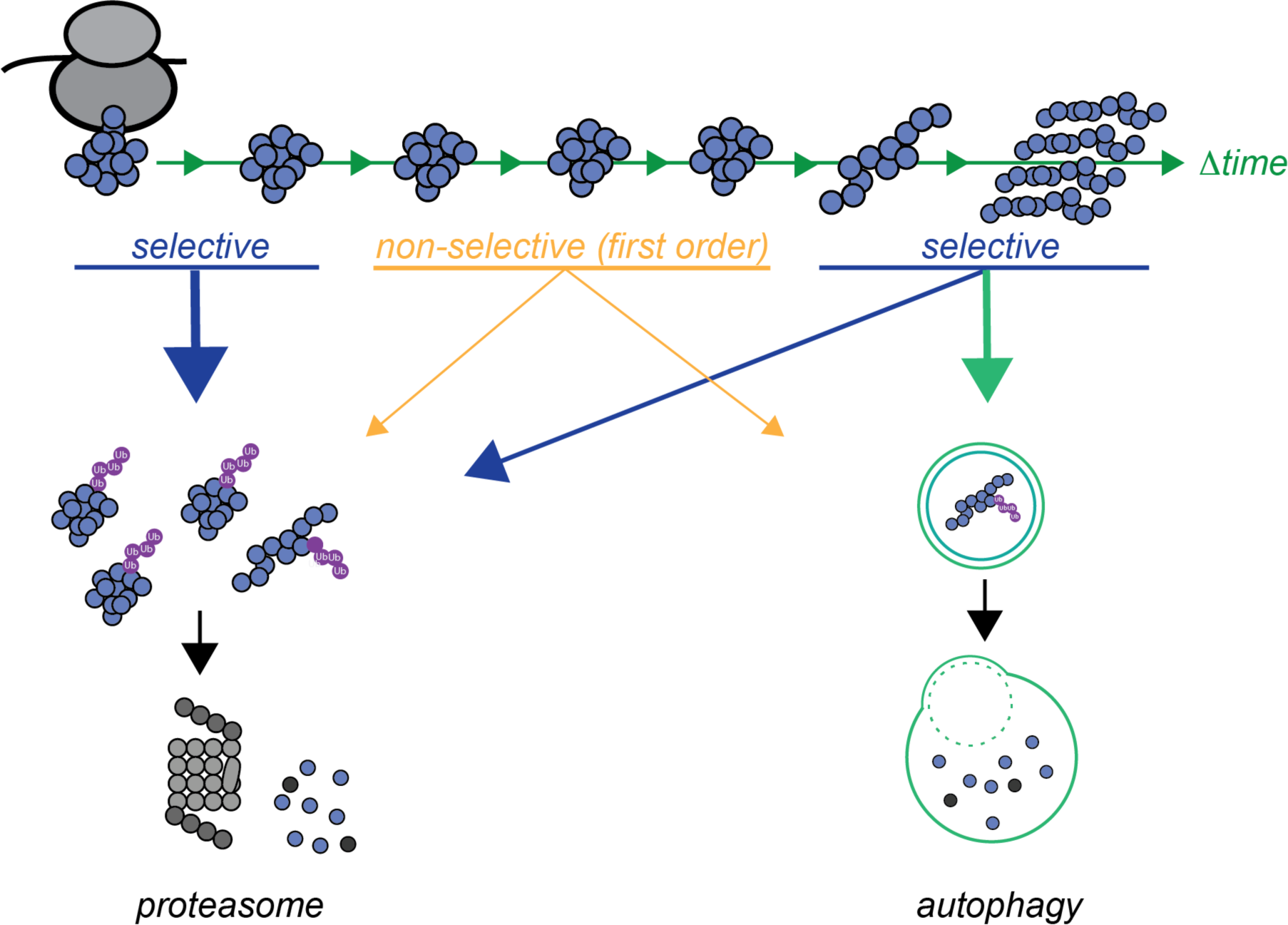
A model for age-selectivity of protein degradation. Significant fractions of proteins are selectively targeted for degradation either immediately after synthesis or after residing in the cell for extended periods of time. These two populations compose relatively small fractions of a protein’s steady-state population, which is largely degraded in a non-selective manner and follows first-order kinetics. Selective degradation of young proteins constitutes a greater fraction of proteasomal flux (indicated by the larger arrow). Older proteins are ubiquitinated and degraded at relatively slower rates and can be targeted to non-proteasomal, yet still selective degradation pathways such as autophagy.

This general model addresses an apparent paradox in the field of protein turnover that has been recognized since the 1960s (43–45). Ostensibly, the idea of selective protein degradation seems to contradict the fact that protein degradation can usually be modelled based on first order exponential kinetics. Indeed, the notion that a protein has a single distinct degradation rate constant and half-life implies that its clearance is a non-selective random process where every molecule in the population has the same chance of degradation regardless of its age. However, mechanistic studies of protein degradation pathways such as UPS and selective autophagy leave little doubt that a cell’s protein quality control mechanisms can specifically target and clear nonfunctional proteins at specific stages of their life cycle. The proposed model avoids this contradiction because the selectively targeted portions of the protein population (damaged young and old proteins) are not expected to constitute a significant fraction of the accumulated proteome at steady state. Thus, in continuous labeling experiments, turnover of cellular proteins may give the appearance of non-selective first order kinetics even though portions of their populations are selectively cleared at faster rates.

Whether the kinetic trends uncovered here for cultured human fibroblasts are representative of other cell types and tissues *in vivo* remains to be determined. Proteome birthdating may prove helpful in such studies and, more generally, may allow for analyses of diverse post translational modifications and cellular processes that discriminate among proteins on the basis of age.

## Materials and methods

### Cell Culture and Stable Isotope Labeling

Human dermal fibroblasts (HCA2-hTert) were cultured in Eagle’s Minimum Essential Medium (ATCC) supplemented with 15% fetal bovine serum (Thermo Fisher) and 100 µg/ml Primocin (Invitrogen) at 37 °C with 5% CO_2_ and 4% O_2_. Cells were adapted to labeling conditions by growth in this media supplemented with 15% dialyzed fetal bovine serum (Thermo Fisher) and 100 ug/mL Primocin (Invitrogen). Quiescent cells were gradually adapted to this media over a period of 4 days. Cells were then split into experimental plates and allowed to achieve a state of quiescence by contact inhibition as described previously (28).

For dynamic SILAC experiments, a series of plates, representing each time point, were switched to labeling media containing MEM for SILAC (Thermo Scientific) supplemented with dialyzed fetal bovine serum (Thermo Scientific), Primocin, and l-arginine:HCl (13C6, 99%; Cambridge Isotope Laboratories) and l-lysine:2HCl (13C6, 99%; Cambridge Isotope Laboratories) at concentrations of 0.13 g/L and 0.087 g/L, respectively. Cells were collected after 0, 24 h, 72 h, 120 h, 168 h, and 336 h of labeling. Cells were removed from plates by trypsinization, washed with PBS buffer, and flash frozen as pellets prior to further analysis.

For proteome birthdating experiments, adapted fibroblasts were plated on one plate per replicate and allowed to become quiescent. Labeling media containing K_202_, K_040_, K_602_, and K_080_ (Thermo Scientific and Cambridge Isotope Laboratories) were made as above. Cells were labeled according to one of the two following labeling schemes as described in Results: 4 d K_202_, 2 d K_040_, 1 d K_602_, and 6 h K_080_; 5 d K_202_, 3 d K_040_, 1 d K_602_, and 6 h K_080_. Prior to switching of labels at each timepoint, cells were washed thoroughly with PBS three times. At the end of the time-course. Cells were processed and stored as above.

### Proteasome Inhibition

To achieve proteasomal inhibition in unlabeled cells, 5mg of MG132 (EMD Millipore) was resuspended in 1 mL of DMSO (Corning) and cultures were treated at 0, 5, 10 or 25 µm for 6 hours. To inhibit the proteasome in birthdated cells, 10 µm MG132 was added to cells coincident with the addition of the K_080_ label (6 h prior to the end of the time course). All cells were trypsinized, washed with PBS, and flash frozen as cell pellets prior to further analysis.

### Cell Lysis and Protein Digestion

Cell pellets were lysed in RIPA buffer (EMD Millipore) supplemented with EDTA Free Protease and Phosphatase Inhibitors (Thermo Scientific). Following incubation with rotation for 30 minutes, lysates were centrifuged for 15 minutes at 19,000 xg at 4 °C. Protein concentrations of the cleared lysates were measured by BCA assays using a plate reader (Thermo Scientific). 25 *µ*g of each lysate was brought up to 25 *µ*l in 50 mM TEAB buffer (Thermo Scientific). Disulfide bonds were reduced with the addition of DTT (Thermo Scientific) to a final concentration of 2 mm with incubation at 55 °C for 1 h. Reduced cysteines were alkylated by the addition of iodoacetamide (Thermo Scientific) at a final concentration of 10 mM by incubating in the dark at room temperature for 30 minutes. Subsequently, 12% phosphoric acid was added to a final concentration of 1.2% and samples were diluted 6-fold in 90% methanol in 100 mM TEAB. Samples were then loaded onto S-TRAP columns (ProtiFi), washed with 90% methanol in 100mM TEAB twice. 20 *µ*L solutions containing 1 *µ*g trypsin (Thermo Scientific) or 1 *µ*g Lys-C (Thermo Scientific) were added to the columns and incubated at 37 °C overnight. Samples were then eluted in 40 *µ*L of 0.1% TFA in H_2_O and then 40 *µ*L of 50/50 ACN:0.1% TFA and subsequently pooled together. Peptides were then dried down and brought up in 100 mM ammonium hydroxide for high pH fractionation.

### High pH Fractionation

Digested samples were reconstituted in 50 μl of 100 mM ammonium hydroxide and fractionated using MagicC18 Stage Tips. Briefly, columns were prepared by twice adding 50 μl of acetonitrile and centrifuging at 2000 ×g for 2 min, then twice washing with 50 μl 10 mM ammonium hydroxide. Samples were then loaded onto the column and centrifuged at 2000 ×g for 2 min, then washed once with 10 mM ammonium hydroxide. Peptides were subsequently eluted in 16 fractions, with 2%, 3.5%, 5%, 6.5%, 8%, 9.5%, 11%, 12.5%, 14.5%, 15.5%, 17%, 18.5%, 20%, 21.5%, 27%, and 50% acetonitrile in 10 mM ammonium hydroxide. 8 mixed fractions were generated by combining pairs of fractions (2% and 12.5%, 3.5% and 14.5%, etc.) resulting in 8 total mixed fractions. Fractions were dried down by speed vac, resuspended in 12.5 μl of 0.1% TFA, and analyzed as described below.

### Kε-GG peptide immunoprecipitation

Kε-GG peptides were enriched using the PTMScan HS Ubiquitin/SUMO Remnant Motif (Kε-GG) Kit from Cell Signaling Technologies using the provided reagents and protocol. Briefly, cell pellets were lysed in 20 mM HEPES buffer containing 9 M urea, which also inhibited residual de-ubiquitinating enzyme activity. Cystines were reduced with 5mM DTT for 1 hour at room temperature and alkylated by 50 mM IAA for 15 minutes at room temperature in the dark. Subsequently, samples were diluted to <2M urea with 20 mM HEPES pH 8.0 and digested with trypsin at a ratio of 1mg trypsin to 37.5 mg lysate at 37 °C overnight. Total digested peptides were recovered by Sep-Pak C18 columns (Waters). Digested Kε-GG peptides were then enriched using the selective monoclonal antibodies conjugated to magnetic beads as provided in the kit. Purified Kε-GG peptides were eluted in 0.15% TFA, desalted with MagicC18 Stage Tips, and then analyzed by LC-MS/MS.

### Western Blots

Cell pellets were prepared and stored as described above and subsequently lysed in RIPA buffer (EMD Millipore) while being rotated for 30 min at 4 °C. Lysates were then cleared by centrifugation at 19,000 xg for 15 minutes at 4 °C. BCA assay was then performed to determine protein concentrations (Thermo Scientific). Lysates were mixed with sample buffer (Bio-Rad), boiled, and centrifuged at 19,000 xg for 5 min. 22.5 *µ*g of each sample was then loaded onto 4-20% Bio-Rad precast gels (Bio-Rad).

Electrophoresis was carried out for 35 minutes at 250 V using a Powerpac HV power supply (Bio-Rad). Some gels were then stained with Coomassie G250 Biosafe Stain. For western blots, replicate gels were transferred to Trans-Blot Turbo Midi PVDF membranes (Bio-Rad) with the Trans-Blot SD Semi-Dry Electrophoretic Transfer Cell (Bio-Rad). Blots were washed for 5 min with TBST, then blocked with LI-Cor Intercept Blocking Buffer for 30 minutes (LI-COR Biosciences). Blots were incubated with primary anti-ubiquitin antibody (E4I2J Cell Signaling Technologies) in blocking buffer overnight. Blots were washed for 5 min 3 times at room temperature with TBST, then secondary antibody (Dylight 800 secondary anti-rabbit, Thermo Scientific) in blocking buffer for 1 hour at room temperature. Blots were washed again 3 times for 5 minutes in TBST and finally stored in TBS. Images were acquired with Bio-Rad ChemiDoc MP Imaging System (Bio-Rad) and processed using ImageJ.

### LC-MS/MS Analysis

Peptides from each fraction were injected onto a homemade 30 cm C18 column with 1.8 μm beads (Sepax), with an Easy nLC-1200 HPLC (Thermo Fisher), connected to a Fusion Lumos Tribrid mass spectrometer (Thermo Fisher). Solvent A was 0.1% formic acid in water while solvent B was 0.1% formic acid in 80% acetonitrile. Ions were introduced to the mass spectrometer using a Nanospray Flex source operating at 2 kV.

For dSILAC experiments, the gradient began at 3% B and held for 2 minutes, increasing to 10% B over 7 minutes, increased to 38% over 68 minutes, ramped up to 90% over 3 minutes and held for 3 minutes before ramping down to 0% over 2 minutes and re-equilibrating the column for 7 minutes for a total run time of 90 minutes. The Fusion Lumos was operated in data-dependent mode, with MS1 and MS2 scans acquired in the Orbitrap (OT) and ion trap (IT) respectively. The cycle time was set to 1.5 seconds, Advanced Peak Determination (ADP) was set to “TRUE” and Monoisotopic Precursor Selection (MIPS) was set to “Peptide.” MS1 scans were performed over a range of 375-1400 m/z with a resolution of 120K and m/z of 200, an automatic gain control (AGC) target of 4e5, and a maximum injection time of 50 ms. Peptides with a charge state between 2-5 were chosen for fragmentation. Precursor ions were fragmented by collision-induced dissociation (CID) using a collision energy of 30%, activation time of 25 ms and activation Q of 0.25 with an isolation width of 1.1 m/z. The IT scan rate was set to “Rapid” with a maximum injection time of 35 ms and an AGC target of 1e4. Dynamic exclusion was set to 20 seconds and to exclude after 1 time with both low and high mass tolerance set to 10 ppm with exclude isotopes set to “TRUE.”

For the NeuCode experiments in Figures 2 and Kε-GG Ips, the gradient began at 3% B and held for 2 minutes, increasing to 10% B over 6 minutes, increased to 48% over 95 minutes, ramped up to 90% over 5 minutes and held for 3 minutes before returning to starting conditions over 2 minutes and re-equilibrating the column for 7 minutes for a total run time of 120 minutes. The Fusion Lumos was operated in data-dependent mode, with MS1 and MS2 scans acquired in the OT and IT respectively. The cycle time was set to 2 seconds, ADP was set to “TRUE,” and MIPS was set to “Peptide.” MS1 scans were performed over a range of 350-1100 m/z with a resolution of 500K at m/z of 200, an AGC target of 1e6, and a maximum injection time of 50 ms. Peptides with a charge state between 2-6 were chosen for fragmentation. Precursor ions were fragmented by higher energy collision dissociation (HCD) using a collision energy of 30% with an isolation width of 1.1 m/z. The IT scan rate was set to “Turbo” with a maximum ion injection time of 15 ms and an AGC target of 2e3. MS2 scans were performed over a range of 200-1200 m/z. The minimum intensity threshold was set to 5e3 and maximum intensity threshold was set to 1e20. Dynamic exclusion was set to 5 seconds and to exclude after 1 time with a low mass tolerance of 0.55 m/z and a high mass tolerance of 1.55 m/z with exclude isotopes set to “TRUE.”

For the NeuCode experiments on the unmodified peptides in Figure 3, data the gradient and data acquisition were performed identically with the following exceptions. Two separate MS1 scans were performed, one at low resolution (30K) for precursor fragmentation, and the other at high resolution (500K) for NeuCode quantification. The 30K MS1 scan was performed over a range of 350-1100 m/z while the 500K MS1 scan was performed over a range of 349-1099 m/z. Both 30K and 500K MS1 scans had a maximum injection time of 100 ms. There were no intensity thresholds set.

For the unlabeled Kε-GG IP experiments in Figure 4, the gradient began at 3% B and held for 2 minutes, increasing to 10% B over 6 minutes, increased to 38% over 95 minutes, ramped up to 90% over 5 minutes and held for 3 minutes before returning to starting conditions over 2 minutes and re-equilibrating the column for 7 minutes for a total run time of 120 minutes. The Fusion Lumos was operated in data-dependent mode, with MS1 and MS2 scans acquired in the OT. The cycle time was set to 2 seconds, ADP was set to “TRUE”, and MIPS was set to “Peptide.” MS1 scans were performed over a range of 375-1400 m/z with a resolution of 120K at m/z of 200, an AGC target of 4e5, and a maximum injection time of 50 ms. Peptides with a charge state between 2-5 were chosen for fragmentation. Precursor ions were fragmented by HCD using a collision energy of 30% with an isolation width of 1.5 m/z. MS2 scans were performed over a range of 200-1200 m/z with a resolution of 15K at m/z of 200, an AGC target of 5e4, and a maximum injection time of 22 ms with maximum injection time type set to “Dynamic.” The minimum intensity threshold was set to 8e4 and maximum intensity threshold was set to 1e20. Dynamic exclusion was set to 30 seconds and to exclude after 1 time with low and high mass tolerances of 10 ppm and exclude isotopes set to “TRUE.”

### Database Searches

Except for dSILAC experiments (see below), all raw files were searched using SEQUEST within the Proteome Discoverer software platform, version 2.4 (Thermo Scientific) against the Uniprot human database (11/18/2019, 20,541 sequences). For high pH fractionation experiments, the fractions were combined within Proteome Discoverer to create one report. Trypsin was selected as the enzyme, allowing up to two missed cleavages. MS1 tolerance was set to 10 ppm, while the MS2 tolerance was set to 0.6 Da. Due to “Indistinguishable Channel” errors within Proteome Discoverer when trying to differentiate between similar isotopes, two database searches were required for each file. The first search included oxidized methionine, Met-loss on protein N-term, NeuCode 202 (+4.0008 Da), and NeuCode 602 (+8.0142 Da) on lysine as variable modifications, while carbamidomethyl on cysteine was selected as a fixed modification. The second search replaced NeuCode 202 and 602 with NeuCode 040 (+4.0251 Da) and NeuCode 080 (+8.0502 Da) as variable modifications, with all other parameters remaining unchanged.

For KGG-enriched samples, custom modifications needed to be created due to GG (+114.0429 Da) and NeuCode labels occurring on lysine at the same time. Once again, two searches per file were done, with GG + NeuCode 202 (+118.0437 Da) and GG + NeuCode 602 (+122.0571 Da) included in the first search, and GG + NeuCode 040 (+118.0680 Da) and GG+ NeuCode 080 (+122.0931 Da) included in the second search. All other parameters listed above remained the same, except that the number of missed cleavages was increased to 3.

For all analysis, Percolator was used as the FDR calculator, filtering out peptides which had a q-value greater than 0.01.

For dSILAC experiments, all raw files were searched using FragPipe platform that includes MSFragger v20.0 against the Uniprot human database (05/03/2022, supplemented with false decoys and contaminants via FragPipe). As SILAC experiments were not fractionated prior to LC-MS/MS, we submitted 6 samples from cells that were differentially labeled with heavy lysine (13C6) and arginine (13C6) for either 0d, 1d, 3d, 5d, 7d, 14d. Strict trypsin was selected as the enzyme, allowing up to two missed cleavages. MS1 tolerance was set to 20 ppm, while the MS2 tolerance was set to 0.6 Da. The search included the appropriate variable modifications of oxidized methionine, Met-loss on protein N-term and accompanying acetylation, and heavy labeled Lysine and Arginine (+6.020129 Da), while carbamidomethyl on cysteine was selected as a fixed modification. Global protein half-lives were then calculated by calculating the L/H+L ratio of Lysine and Arginine for each peptide/protein at each time point and then generating kinetic curves for the ratios over the 14-day period.

### Measurement of peptide and protein kdeg, half-lives and ages

The kinetic model and method for determining first order degradation rate constants (*k_deg_*) from dSILAC experiments has been described in detail previously (28). Briefly, heavy to light (H/L) SILAC ratios were determined at the peptide level using the software FragPipe (46). The H/L ratios for each peptide were converted to fraction labeled (H/(H+L)) measurements. To determine *k_deg_* at the protein level, fraction labeled measurements for all peptides and time-points mapped to specific proteins were combined in single aggregated kinetic plots. Protein degradation was modeled as a first order reaction and as cells were quiescent, the rate of dilution due to cell division was assumed to be negligible. The kinetic plots were fitted to the following single exponential function using non-linear curve fitting and the Levenberg-Marquardt algorithm:

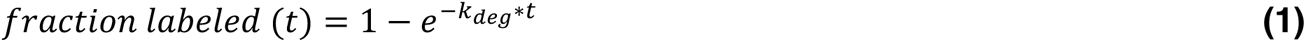

Where t is the continuous labelling time. *K_deg_* measurements were converted to protein half-lives using the following equation:

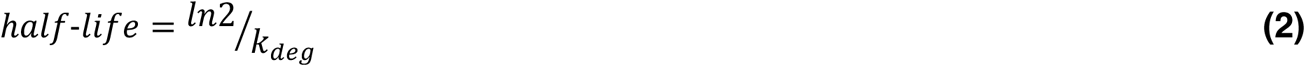

For determining *k_deg_* from proteome birthdating data, database searches were conducted by Proteome Discoverer (PD) as described above and MS1 spectra were exported as .ms1 files using msConvert (47). Intensities of PSMs labeled with K_000_, K_202_, K_040_, K_602_,and K_080_ were quantified from .ms1 files using a Mathematica notebook written in house. For each PSM, intensities of the five labeled forms were converted to relative fractional populations. For analyses conducted at peptide and protein levels, all corresponding PSMs were aggregated in single kinetic plots. In accordance with a first order kinetic model, the fractional population of each label for each PSM is related to *k_deg_* by the following equation:

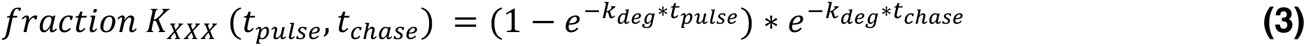

Where *t_pulse_* and *t_chase_* are the distinct pulse and chase times for each label. For example, in a time course where initially unlabeled (K_000_) cells are sequentially labeled with K_202_, K_040_, K_602_ and K_080_ for 4 d, 2 d, 1 d and 0.25 d, *t_pulse_* for the five labels are ∞ d, 4 d, 2 d, 1 d and 0.25 d, and *t_chase_* are 7.25 d, 3.25 d, 1.25 d, 0.25 d and 0 d, respectively. *K_deg_* values were determined using non-linear curve fitting of aggregated birthdating data to equation 3 employing the Levenberg-Marquardt algorithm. Goodness-of-fit for the measured parameters were determined by t-statistic and R^2^ measurements. To pass quality control thresholds, *k_deg_* measurements were required to fulfill all three of the following criteria: 1) at least two unique PSMs were quantified, 2) t-statistic for the nonlinear fit exceeded 3.0, and 3) R^2^ for the nonlinear fit exceeded 0.9.

*k_deg_* measurements determined by proteome birthdating were converted to half-lives using equation 2. As proteins were assumed to obey first order decay kinetics, half-lives could be equated to the “median age” of the steady-state protein population. Hence, at steady state, half of a given protein’s population were assumed to be older than its half-life/median age, and half were assumed to be younger than its half-life/median age.

### Gene ontology enrichment analysis

Mappings of GO terms to UniProt accessions, and their relational hierarchies were obtained from the gene ontology database (downloaded June 2023). For all proteins mapped to a given GO term, measured parameters (e.g. half-lives in Figure S2 or Log_2_ age ratio between Kε-GG and non-Kε-GG peptides) were collected. The p-values for differences between the distribution of these values for a given GO term in comparison to the entire measured dataset were determined using the Mann-Whitney U test.

## Author Information

### Author Contributions

MM, KW, AS, VG, AB, & SG conceived of the experiments. MM, SB, MH, FA, KW, KS, JH, JM, SAB performed the experiments. MM, KW, KS, & SG analyzed data and conducted bioinformatic analyses. MM, KW, KS, & SG wrote the manuscript.

### Notes

The authors declare no competing financial interest.

## Acknowledgements

The authors would like to thank the members of the Ghaemmaghami, Gorbunova, Seluanov, and Fu laboratories for helpful discussions.

## Data Availability

All raw and processed data are available in the included Supporting Information and at the ProteomeXchange Consortium via the PRIDE partner repository (accession number PXD045886). Currently, the data can be accessed with the username and password LH2fjiKd.

## Abbreviations

ACN: acetonitrile
AGC: automatic gain control
ALDH: aldehyde dehydrogenase
APD: advanced peak determination
CID: collision induced dissociation
CPN60: chaperonin 60
DriP: Defective Ribosomal Products
DMSO: dimethyl sulfoxide
DNPK1: DNA-dependent protein kinase catalytic subunit
dSILAC: dynamic stable isotope labelling by amino acids in cell culture
DTT: dithiothreitol
ER: endoplasmic reticulum
ERAD: endoplasmic reticulum associated degradation
FDR: false discovery rate
GO: gene ontology
HCD: higher energy collision dissociation
HEPES: 4-(2-hydroxyethyl)-1-piperazineethanesulfonic acid
HPLC: high pressure liquid chromatography
hTERT: human telomerase
IAA: iodoacetamide
IP: immunoprecipitation
IT: ion trap
LC-MS/MS: liquid chromatography-tandem mass spectrometry
MIPS: monoisotopic precursor selection
MS: mass spectrometry
MW: molecular weight
NeuCode: neutron encoded
nLC: nanoLiquid chromatography
OT: Orbitrap
PD: proteome discoverer
PSM: peptide spectral match
PTM: post-translational modification
PVDF: Polyvinylidene fluoride
RIPA: Radioimmunoprecipitation Assay
ROS: reactive oxygen species
RQC: ribosomal quality control
RT: retention time
SD: semi-dry
SRP: signal recognition particle
TEAB: triethylammonium bicarbonate
TFA: trifluoroacetic acid
UPS: ubiquitin proteasome system

## Supplementary Figures

**Figure S1.**
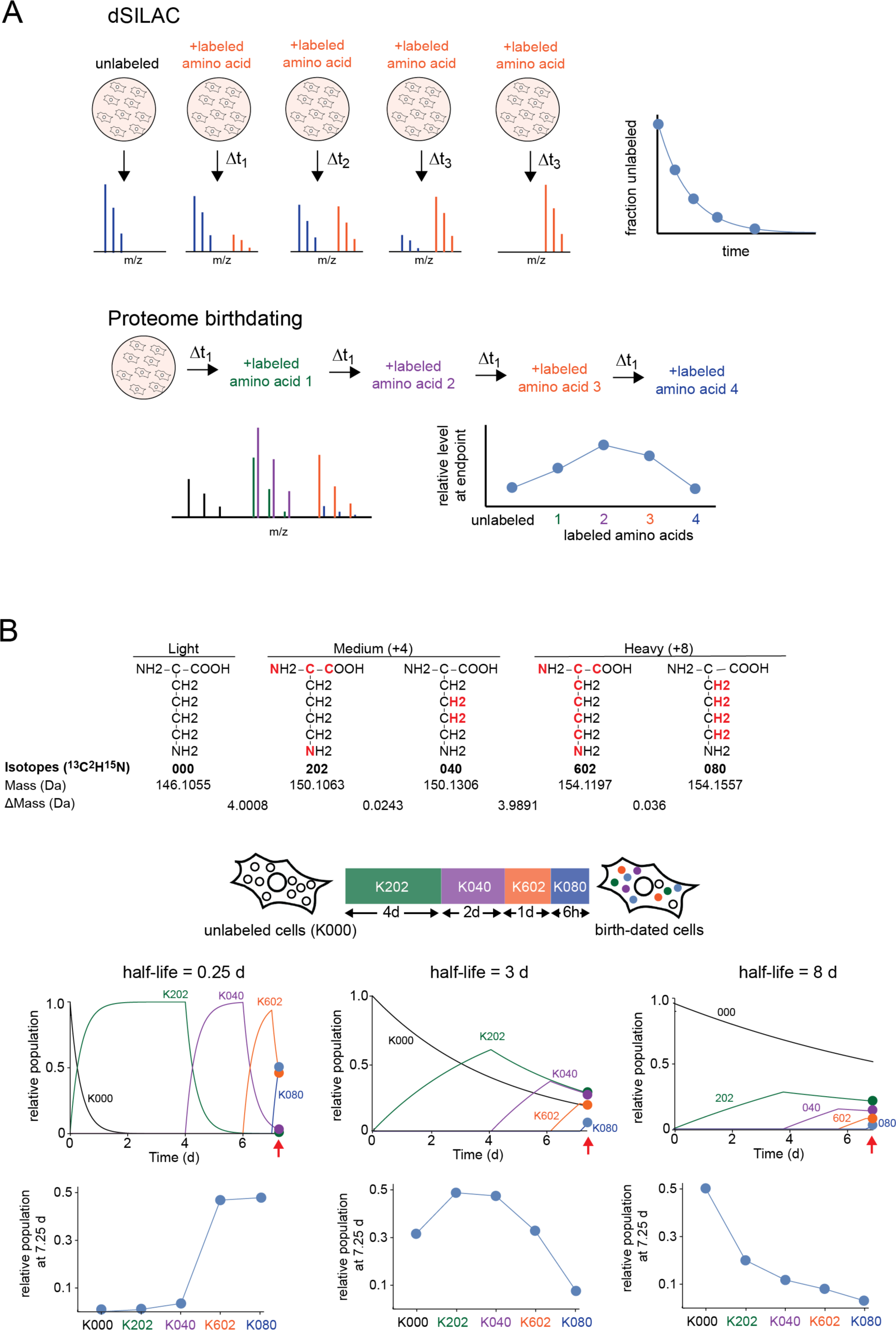
Comparison of dSILAC and proteome birthdating as methods for measurement of global protein half-lives (Related to Figure 1) **A)** General experimental designs of dSILAC and proteome birthdating. In dSILAC, a single isotopically labeled amino acid is added to several distinct biological samples that constitute each timepoint. Protein turnover kinetics are then analyzed by measuring changes in fractional labeling of proteins as a function of time. In proteome birthdating, a number of different isotopically labeled amino acids are added and removed from the same biological sample. Relative levels of differentially labeled variants of proteins are then measured at the endpoint of the experiment. **B)** Structures show the five isotopic variants of lysine used in this paper. The red atoms indicate the sites of heavy isotope incorporation. ΔMass refers to the difference in mass between adjacent variants. The kinetic plots show theoretical changes in relative populations of K_XYZ_-labeled peptides over time during the labeling time course described in Figure 1A. The bottom plots show the relative populations of K_XYZ_-labeled peptides at the end of the time course. Kinetic and endpoint plots are shown for three theoretical peptides with different half-lives as described in Figure 1B.

**Figure S2.**
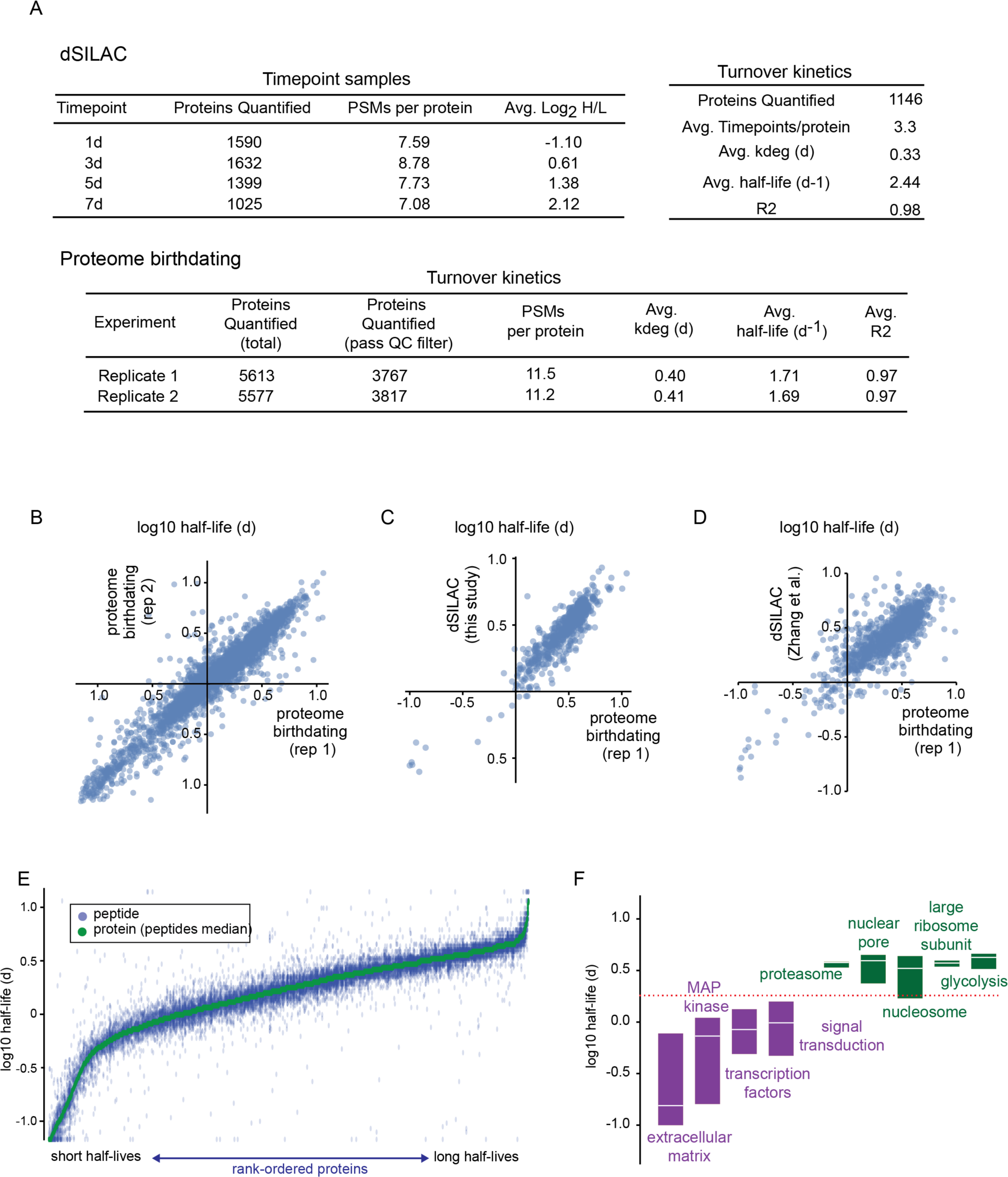
Coverage and accuracy of protein half-life measurement determined by proteome birthdating (Related to Figure 2) **A)** Coverage statistics of dSILAC and proteome birthdating experiments. Details of the analyses and quality control (QC) filters are described in Materials and Methods. Datasets of the measured parameters are tabulated in Supplementary Table S1. **B)** Pairwise comparison of protein half-lives measured in two replicate proteome birthdating experiments. **C)** Pairwise comparison of protein half-lives measured in with proteome birthdating (replicate 1) and dSILAC performed in this study. **D)** Pairwise comparison of protein half-lives measured in with proteome birthdating (replicate 1) and by dSILAC in Zhang et al. (28). **E)** Rank-size distribution plots showing half-lives of peptides measured by proteome birthdating (replicate 1). Vertical columns of blue points on the plot represent data for peptides matched to specific proteins. The green points are the median of peptide-level half-life measurements for each protein. Proteins are rank ordered based on their half-lives. Note that the range of measured half-lives for peptides within each protein is narrow relative to the entire range of all measured peptides within the proteome. **F)** Half-lives of peptides measured by proteome birthdating for proteins mapped to the indicated GO terms. The plotted GO terms are a subset of terms whose constituent proteins have half-lives are significantly longer or shorter than the proteome at large. Box plots indicate the interquartile range of measurements and the line indicates the median. The complete list of GO terms with half-lives that are significantly different than the complete proteome is listed in Supplementary Table S1.

**Figure S3.**
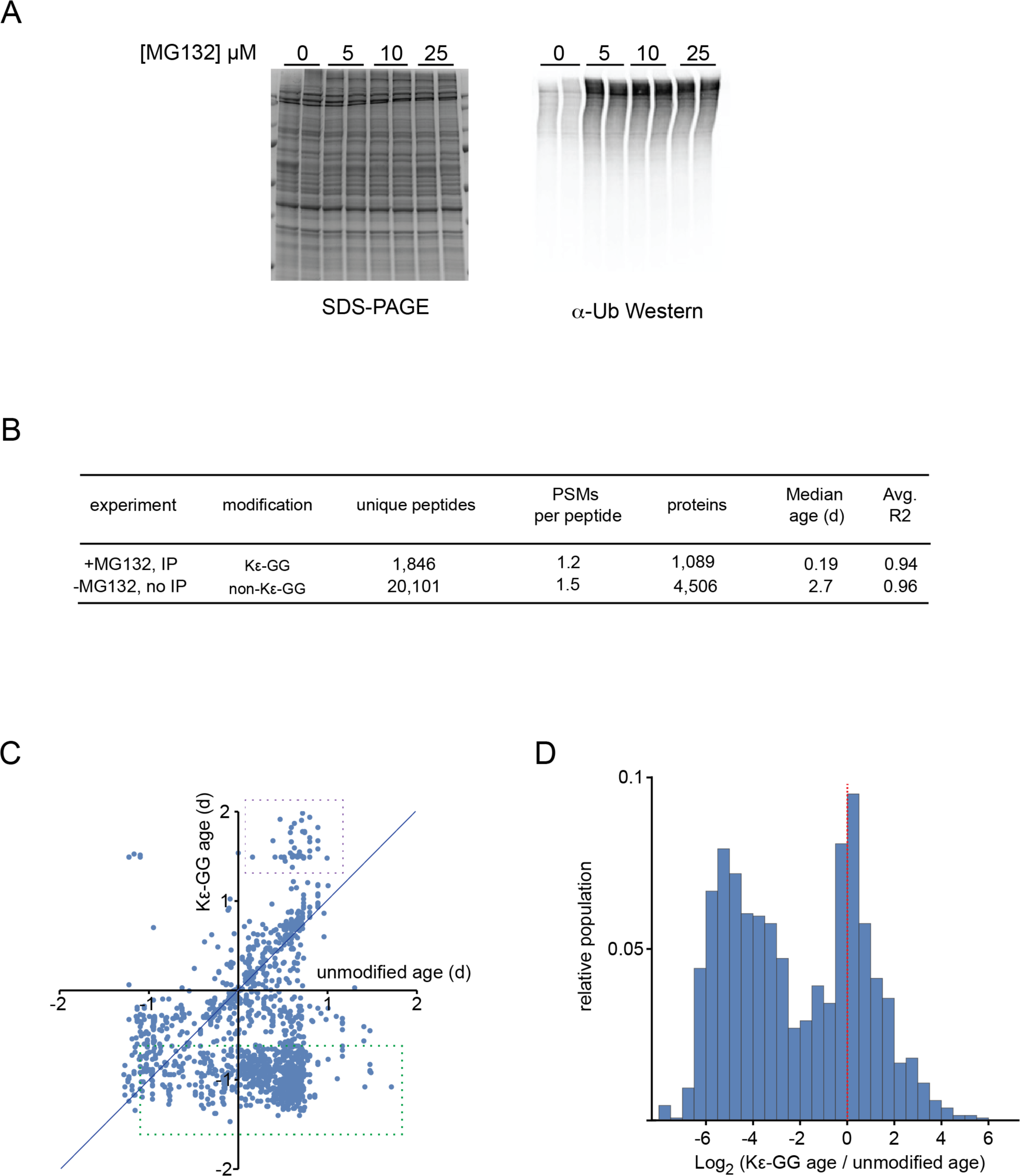
Analysis of the age distribution of Kε-GG peptides by proteome birthdating (Related to Figure 3) **A)** SDS-PAGE and anti-ubiquitin western blots showing the accumulation of ubiquitinated proteins in human fibroblasts after addition of MG132 at different concentrations for 6 hours. Each concentration point was analyzed in duplicate experiments. **B)** Coverage statistics of proteome birthdating experiments for Kε-GG and unmodified peptides. Details of the analyses are described in Materials and Methods. Datasets of the measured parameters are tabulated in Supplementary Table S2. **C)** Pairwise comparison of measured median ages of Kε-GG and unmodified peptides mapped to the same proteins. Green and purple dashed boxes highlight Kε-GG peptides that are, respectively, younger or older than their unmodified counterparts mapped to the same proteins. **D)** Log_2_ ratios of ages of Kε-GG and unmodified peptides mapped to the same proteins.

**Figure S4.**
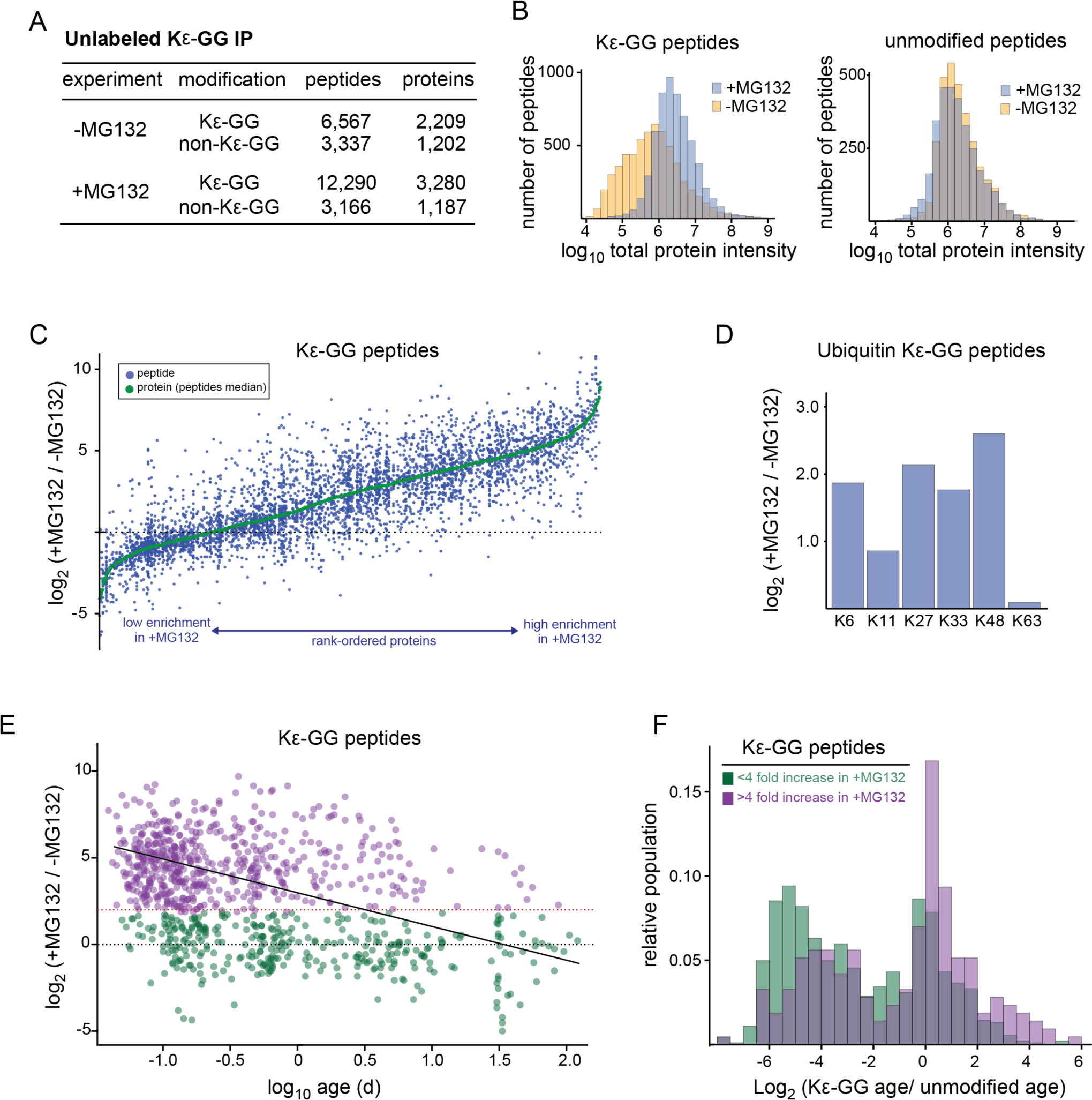
Correlation between proteasomal flux and protein age determined by proteome birthdating (Related to Figure 4) **A)** Coverage statistics for proteome-wide changes in Kε-GG peptide levels upon proteasome inhibition. Datasets of the measured parameters are tabulated in Supplementary Table S3. **B)** Intensity distributions of Kε-GG and unmodified peptides in the presence and absence of MG132. The plots indicate that Kε-GG peptides, but not unmodified peptides, increase their levels upon proteasomal inhibition. **C)** Rank-size distribution plots showing Log_2_ fold increases in levels of Kε-GG peptides upon proteasomal inhibition. Vertical columns of blue points on the plot represent data for peptides matched to specific proteins. The green points are median peptide-level measurements for each protein. Proteins are rank ordered based on median peptide measurements. Note that MG132-induced changes in levels of Kε-GG peptides mapped to the same proteins are highly variable. **D)** Log_2_ fold increases in levels of Kε-GG peptides mapped to ubiquitin upon proteasomal inhibition. **E)** Correlation between MG132-induced changes in levels of Kε-GG peptides and their age. **F)** Log_2_ ratios of ages of Kε-GG and unmodified peptides mapped to the same proteins. In E and F, peptides whose levels increased by greater or less than a factor of four are shown in purple and green colors, respectively. The analysis in E and F indicate that, in general, younger Kε-GG peptides accumulate more in the presence of MG132 in comparison to older Kε-GG peptides.

